# Integrating Genomic Annotations and Traits Dependencies for single-nucleotide polymorphisms Prioritization with Causal Concept Bottleneck Models

**DOI:** 10.64898/2026.09.04.749112

**Authors:** Francesco De Santis, Daniele Malpetti, Francesco Gualdi, Francesca Mangili

## Abstract

Predicting common traits from single-nucleotide polymorphism (SNPs) data is challenging due to polygenicity, small effect sizes, and the presence of potentially mediated or spurious cross-trait associations. We propose a modeling approach that combines genomic annotations with known cross-trait relations by leveraging Causally Reliable Concept Bottleneck Models (C^2^BM), a deep learning architecture that factors the joint trait distribution over a graph of interpretable concepts. This design allows trait predictions to leverage information from other observed traits in addition to genomic inputs. Furthermore, the interpretable architecture of the model enables us to investigate how specific trait–trait relationships influence SNP-level predictions. We evaluate the approach on a multi-trait GWAS dataset covering five traits and show that C^2^BM improves predictions when ground-truth labels for related traits are available. Moreover, by analyzing variations in how trait–trait relationships influence predictions, we postulate that such differences may reflect the presence or absence of shared genetic mechanisms or indirect effects.

## 1 Introduction

In recent years, Genome Wide Association Studies (GWAS) have identified many associations between SNPs and traits. However, further work is needed to better characterize the interactions between genetic factors and traits. This is due to the high polygenicity of the disorders, their typically small effect sizes, and the presence of cross-trait associations (pleiotropy). While such associations may reflect shared biological mechanisms, they can also arise through mediated effects or spurious associations induced by confounding factors such as linkage disequilibrium, thereby complicating the identification of direct genetic effects if not explicitly modeled [1].

Over the past decade, several computational methods have been developed to assess the functional impact of genomic SNPs, integrating diverse annotations into predictive scores (e.g., CADD [2]) and, more recently, leveraging deep learning to capture complex, non-linear effects. However, these approaches are often optimized for rare or coding SNPs and show limited performance on common GWAS SNPs, particularly in non-coding regions [3]. Moreover, most methods assign a single, organism-wide score to each SNP, whereas trait-specific models have been shown to improve prioritization [4]. At the same time, GWAS analyses are typically performed on single traits, even though the widespread cross-trait associations observed for complex traits suggest the presence of shared biological pathways or genetic mechanisms across apparently distinct phenotypes [5]. These shared architectures may offer predictive gains when leveraged through multi-trait analyses, but care is required to distinguish true cross-trait signal from confounding factors.

To address these challenges, we leverage Causally Reliable Concept Bottleneck Models (C^2^BM) [6], a concept-based deep learning framework that structures prediction through a causal graph over interpretable concepts. In our setting, this causal structure is used to encode previously established relationships between traits, such as known risk-factor or comorbidity links, e.g., BMI as a risk factor for Coronary Artery Disease.

We apply this framework to model SNP–trait associations using data from the GWAS Catalog, representing SNPs through the same genomic annotations employed by CADD v1.7 [2]. Our approach jointly models associations across multiple traits while remaining trait-specific in its predictions. Furthermore, the C^2^BM architecture provides greater interpretability than conventional deep learning approaches, as SNP effects are mediated through explicit trait relationships. This structure enables the analysis of individual trait–trait dependencies and their influence on SNP–trait association predictions.

## 2 Methodology

We consider high-dimensional genomic data in which each SNP from the GWAS Catalog is represented by a feature vector **x** ∈ ℝ^*d*^ of genomic annotations derived from CADD. In addition, we observe *K* = 5 traits **c** ∈ {0, 1} ^*K*^ (e.g., Crohn’s disease). Our approach consists of two stages, illustrated in Fig. 1: first, we infer a causal graph capturing relationships between traits; second, we use this graph within a C^2^BM framework to model SNP–trait associations.

**Figure 1.**
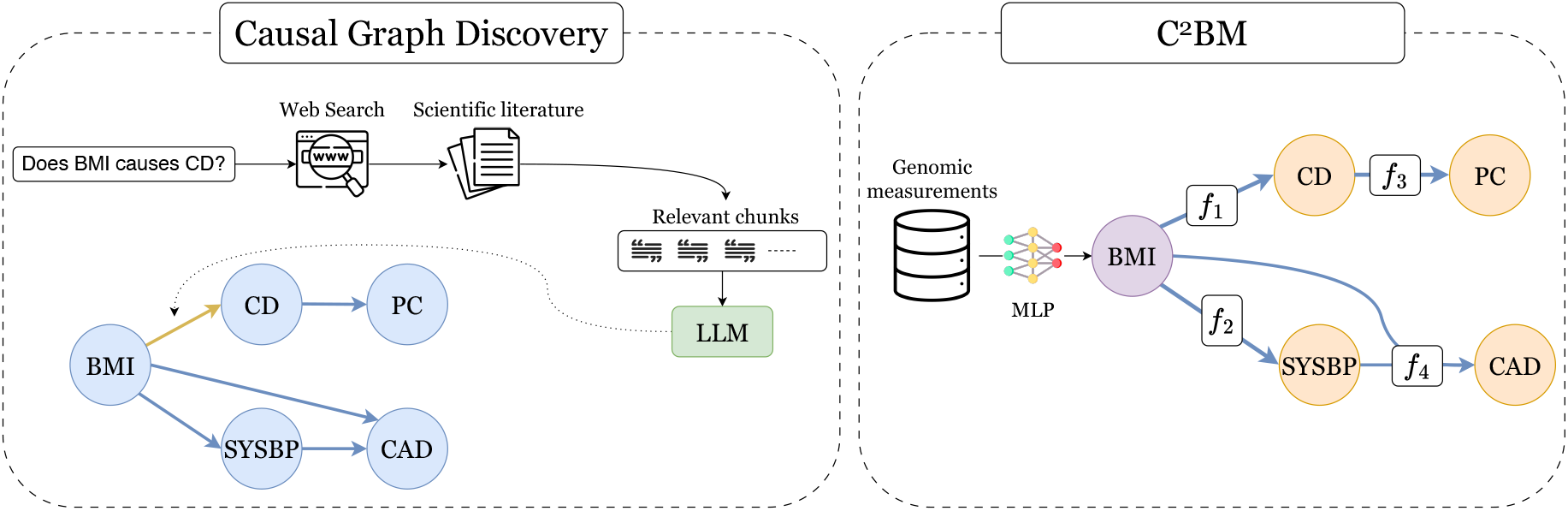
Overview of the proposed approach. **Left:** Causal graph discovery pipeline. **Right:** C^2^BM architecture. Root traits are predicted directly from the genomic input. Non-root traits are predicted via an adaptive linear combination of their causal parents’ predictions and the genomic input.

### Causal graph discovery

Inspired by [6], we estimate a causal graph through a fully automated retrieval-augmented generation (RAG) pipeline. For each ordered pair (*c*_*i*_, *c*_*j*_), GPT-4o generates several search queries (e.g., “Is *c*_*j*_ influenced by *c*_*i*_?”), a web engine retrieves the most relevant pages, and the retrieved content is split into chunks encoded by a pre-trained transformer. The top-8 chunks ranked by cosine similarity to the probe “Does *c*_*i*_ cause *c*_*j*_?” are then passed to GPT-4o, which returns one of three verdicts (*c*_*i*_*→c*_*j*_, *c*_*j*_*→c*_*i*_, or *no relation*) together with a confidence score, defaulting to *no relation* when evidence is absent or contradictory. Pairwise verdicts are assembled into a directed acyclic graph (DAG), with cycles resolved by removing the edge with the lowest confidence score.

### Model architecture

C^2^BM [6] is a recent interpretable model belonging to the broader family of Concept Bottleneck Models (CBMs) [7, 8]. C^2^BM performs predictions over a causal graph whose nodes correspond to human-interpretable variables (e.g., traits). This structure enables the factorization of P(**c** | **x**) as follows:

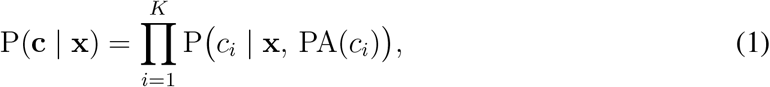

where PA(*c*_*i*_) denotes the parents of *c*_*i*_. Consequently, the prediction for each trait is directly influenced by its parent traits; for example, BMI influences the prediction of CAD. For root traits (i.e., traits with no parents), the probability P(*c*_*i*_ | **x**) is approximated by an MLP operating on genomic measurements, making the prediction equivalent to that of a standard black-box model. For non-root traits, P(*c*_*i*_ | **x**, PA(*c*_*i*_)) is approximated by a learned function:

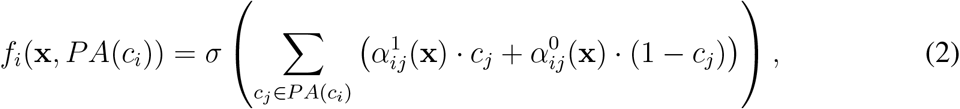

where *σ* is an activation function (e.g., sigmoid for binary *c*_*i*_), and *α*_*ij*_(**x**) denotes MLP-modeled adaptive weights. This allows parent influence to adapt to input **x** while maintaining interpretable parent-child contributions. Causal Graph Discovery

## 3 Experimental settings

We evaluate the proposed approach on a multi-trait genomic dataset, comparing C^2^BM against two baselines in terms of predictive accuracy and ability to leverage inter-trait causal relations.

### Dataset

The dataset is composed of SNPs for **5 different traits**: Systolic Blood Pressure (SYSBP), Body Mass Index (BMI), Coronary Artery Disease (CAD), Prostate Cancer (PC) and Crohn’s Disease (CD), each identified using Experimental Factor Ontology (EFO) terms. For each trait, we selected studies with available summary statistics from the GWAS Catalog [9]. SNPs were considered positively associated with a trait if their p-value was ≤ 5 × 10^™8^. Negative SNPs (p-value *>* 5 × 10^−8^) were randomly sampled to balance the positive cases within each chromosome. Finally, SNPs were annotated using genomic features from CADD v1.7 [2]. Features with more than 40% missing values were removed, while remaining missing values were imputed using mean values for continuous variables and an “undefined” category for categorical variables.

### Experiments and Metrics

We compare C^2^BM against two baselines: a **Bayesian network**, modeling only inter-trait relationships and therefore limited in predictive performance due to the absence of genomic measurements; and a **BlackBox** model, predicting each trait independently from CADD annotations through dedicated MLPs while ignoring inter-trait dependencies. We evaluate all models using the Area Under the ROC Curve (AUC). In addition to this standard metric, we assess how well each model leverages the causal structure among diseases through *interventions* [6, 7]. In an intervention at level *l*, all disease predictions at topological levels 0, …, *l* of the DAG are replaced by their ground-truth labels, and the AUC for the remaining downstream traits is then computed. For the BlackBox, which does not encode inter-trait dependencies, interventions have no effect by construction. In contrast, for both the Bayesian network and C^2^BM, statistically significant improvements in downstream AUC as the intervention level increases would indicate that the causal structure provides informative predictive signal. All experiments are repeated over 10 seeds; for each seed, we randomly resample a 70/10/20 train/validation/test split and reinitialize model weights.

## 4 Results

### Predictive performance

Table 1 reports the AUC obtained by each model under increasing levels of intervention. In the absence of interventions, BlackBox and C^2^BM achieve nearly identical performance across all five traits, indicating that incorporating inter-trait causal structure does not compromise the predictive accuracy attainable from genomic measurements alone. In contrast, the Bayesian network, lacking access to genomic annotations, can only sample from its prior distribution over traits, yielding AUC scores close to 0.500 across all tasks. When ground-truth labels for parent traits are provided, C^2^BM improves predictions for child traits, whereas the BlackBox model remains unchanged by construction. The largest gains are observed for SYSBP, whose AUC increases from 0.769 to 0.795 (+0.026) at intervention level 0, surpassing the BlackBox baseline (0.775), and for CAD, whose AUC increases from 0.781 to 0.806 (+0.025) at intervention level 1, again outperforming the BlackBox baseline (0.786). In both cases, the difference is statistically significant. Notably, the Bayesian network also improves under interventions. These results demonstrate that inter-trait relations carry a predictive signal.

**Table 1:** Per-condition mean AUC (± std. dev.) across different intervention levels, averaged across **10 seeds**.

| Int. Level | Model | BMI | CAD | CD | PC | SYSBP |
| --- | --- | --- | --- | --- | --- | --- |
| No interventions | Bayesian Network | 0.5000±0.0000 | 0.5000±0.0000 | 0.5000±0.0000 | 0.5000±0.0000 | 0.5000±0.0000 |
| | BlackBox | 0.7921±0.0076 | 0.7863±0.0145 | 0.8327±0.0068 | 0.7836±0.0103 | 0.7754±0.0065 |
| | $C^2BM$ | 0.7936±0.0125 | 0.7817±0.0126 | 0.8283±0.0076 | 0.7782±0.0061 | 0.7698±0.0076 |
| Int. at Level 0 | Bayesian Network | – | 0.6131±0.0010 | 0.5797±0.0011 | 0.5667±0.0460 | 0.6373±0.1086 |
| | BlackBox | – | 0.7863±0.0145 | 0.8327±0.0068 | 0.7836±0.0103 | 0.7754±0.0065 |
| | $C^2BM$ | – | 0.7859±0.0124 | 0.8289±0.0078 | 0.7782±0.0062 | 0.7951±0.0060 |
| Int. at Level 1 | Bayesian Network | – | 0.6907±0.0829 | – | 0.5961±0.0251 | – |
| | BlackBox | – | 0.7863±0.0145 | – | 0.7836±0.0103 | – |
| | $C^2BM$ | – | 0.8062±0.0119 | – | 0.7782±0.0062 | – |

### BMI’s effect on CD and SYSBP

To investigate the effect of trait–trait relationships on predictions, we focus on SYSBP and CD, as both have BMI as their only parent in the DAG. This allows us to isolate the contribution of BMI and estimate, for each SNP, the conditional probabilities 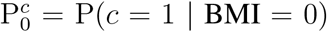 and 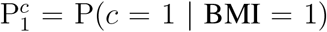. Figure 2 shows the joint distribution of these scores for CD and SYSBP, stratified by SNP association label. For the BMI→CD edge, the joint density of 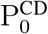 and 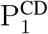 is concentrated along the diagonal, indicating that the model’s prediction for CD is largely insensitive to the BMI parent state. In contrast, for the BMI→SYSBP edge, the density is predominantly shifted above the diagonal, showing that 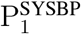 is generally higher than 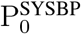. Additionally, both associated and non-associated SNPs exhibit a weak but noticeable concentration in the upper-left region of the plot, corresponding to high values of 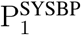 and low values of 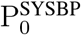.

**Figure 2.**
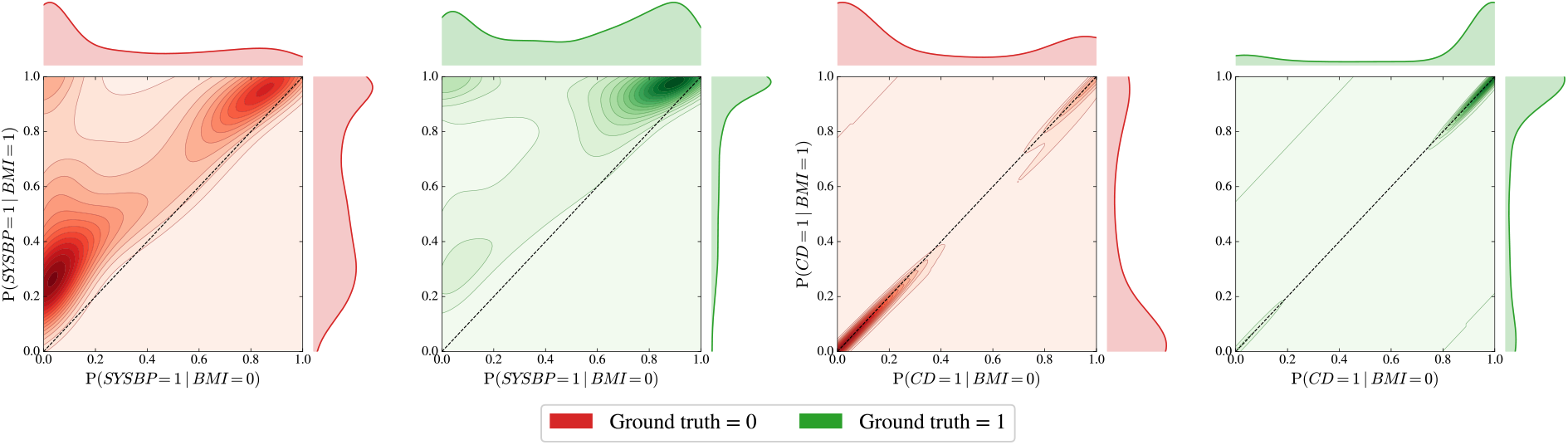
Joint density of P(SYSBP = 1 | BMI = 0) vs. P(SYSBP = 1 | BMI = 1) (two leftmost panels) and P(CD = 1 | BMI = 0) vs. P(CD = 1 | BMI = 1) (two rightmost panels), stratified by ground-truth label. The dashed diagonal indicates equal predictions regardless of the BMI parent state; mass concentrated along it reflects insensitivity to BMI, while off-diagonal shifts indicate that BMI influences the predictions.

## 5 Discussion

In this work, we adopted the C^2^BM framework to jointly model SNP–trait associations by integrating GWAS Catalog data with functional annotations from CADD, while explicitly accounting for known dependencies between traits derived from the scientific literature and represented as a causal network structure.

The SNP-specific scores assigned by the model to each network edge capture shared genetic mechanisms underlying the association of a SNP with both connected traits (pleiotropy). In this sense, the model recognizes features of certain SNPs that suggest a common underlying mechanism, potentially indicative of joint association. Consequently, the observed association with one trait acts as supporting evidence for the presence of such a mechanism, thereby increasing the predicted score for the association with the other connected trait. Our results show that this provides valuable predictive information, particularly when some SNP–trait associations are already known. However, these scores may also reflect spurious pleiotropy, whereby the association of one trait influences the probability of association with a second trait through non-genetic mechanisms.

For instance, BMI is a well-established risk factor for SYSBP, and this relationship may increase the probability of observing SNPs associated with both BMI and SYSBP, even in the absence of a shared genetic basis. While modeling such indirect associations may improve predictive performance, it is important to recognize that they do not correspond to direct genetic effects of the SNP on the target trait. We postulate that these indirect effects, not being driven by SNP-specific genetic mechanisms, would manifest relatively uniformly across SNPs, leading to a systematic increase in P(SYSBP = 1 | BMI = 1). In contrast, true shared genetic mechanisms are expected to be SNP-specific, appearing prominently in only a limited subset of cases while remaining absent for most SNPs. From this perspective, Fig. 2 suggests that neither type of effect is present for BMI and CD, as the vast majority of observations lie along the diagonal. Consistently, no performance improvement is observed in Table 1 when conditioning on BMI.

In contrast, when analyzing BMI and SYSBP, we observe a general upward shift in P(SYSBP = 1 | BMI = 1), suggesting that GWAS Catalog data encode an indirect association between SNPs and SYSBP mediated by BMI, which is captured by the model. This occurs despite the fact that individual studies typically adjust for BMI, potentially indicating that such corrections do not fully remove this source of confounding. In addition, Fig. 2 highlights a smaller subset of SNPs for which the increase in score under the condition BMI = 1 is substantially larger. These cases may reflect shared genetic mechanisms identified by the model and therefore represent promising candidates for further investigation.

## 6 Conclusions

Our results suggest that integrating causal relationships between traits within a concept-based modeling framework can improve the analysis of SNP–trait associations in GWAS data. Beyond predictive performance, the proposed approach provides an interpretable representation of how inter-trait dependencies influence SNP-level predictions, potentially supporting a deeper understanding of the genetic mechanisms underlying complex traits.

This study suffers from some limitations. The dataset contains a large proportion of missing SNP-trait association labels, which makes it harder for the model to accurately estimate the joint distributions between traits and may attenuate the gains attributable to the causal structure. In addition, the interpretation we provide of the SNP- and edge-specific scores assigned by the model is based on a postulated explanation of the underlying mechanisms and thus requires external validation through independent datasets or complementary experimental evidence.

## Acknowledgments

The authors thank Alberto Termine and Zeno Darani for useful discussions.

## Availability of data and software code

The code used to perform the experiments is available at `CBMs-In-Genomics`.

